# Stable maintenance of a hidden switch as a way to increase the gene expression stability

**DOI:** 10.1101/615500

**Authors:** Hiroyuki Kuwahara, Xin Gao

## Abstract

In response to severe stress, wild-type organisms can release alternative phenotypes that are hidden under normal conditions and are associated with underlying genetic variations. A number of such stress-induced phenotypic switchings have been reported to be based on reactivation of hidden thresholds; under the normal condition, a high barrier separating alternative phenotypes ensures the expression of single discrete phenotype, but a severe perturbation can lower the barrier to a level at which to expose cryptic alternatives. While the importance of such threshold-based switches as the mechanism to generate adaptive novelties under variable environments has been appreciated, it still remains elusive how naturally selected organisms can maintain the phenotypic switching capability when such switching has been disused for a long period of time. Here, through the use of computer simulation, we analyzed adaptive evolution of gene circuits under stabilizing selection. We found that different strategies evolved to acquire reduced levels of gene expression noise around the optimum expression level. To incrementally improve the gene expression stability from a founding population with bistable individuals, the evolution consistently took the direction to raise the height of the potential barrier of bistable systems. Our results demonstrate that hidden phenotypic switches can be stably maintained during environmental stasis, facilitating the release of potentially adaptive phenotypic alternatives in the event of substantial perturbations.

## Introduction

The canalization of a discrete phenotype through the buffering of underlying genetic variations is thought to be a general property of naturally selected organisms [1–6]. Substantial perturbations in a wild-type organism can, however, disrupt the normal workings of its genetic developmental system and release alternative phenotypes that would not otherwise be expressed [7–11]. For example, by genetically disrupting the tobacco hornworm, a monophenic species with green larvae, and by applying heat shock, Suzuki and Nijhout were able to derive a polyphenic species that expresses black or green larvae depending on temperature [8]. To release such cryptic phenotypes, wild-type organisms somehow maintain epigenetic switching capabilities in anticipation and lower the threshold of the switch in response to severe perturbations. The stable maintenance of such “hidden” threshold traits in naturally selected organisms suggest that the loss of these obsolete switching mechanisms might somehow incur fitness costs and could be detrimental to the organisms, which could be due, for example, to pleiotropic effects on expressed phenotypes [12]. However, gene sets associated with cryptic phenotypes are likely to be unexpressed under normal conditions and do not seem to contribute to an organism’s fitness. Thus, whereas the presence of phenotypic switching capabilities is commonly presumed in wild-type organisms, it remains elusive exactly how disused switching capabilities to control the expression of alternative phenotypes can be beneficial and maintained in a population for a long period of time [12, 13].

Here, through the use of computer simulation, we report that evolutionary directions towards high levels of gene expression stability under natural selection can explain the stable maintenance of obsolete switching capabilities. We simulated evolution of a gene circuit model that has a potential to exhibit a bistable switch. During the *in silico* evolution, organisms were first placed in a static environment with stabilizing selection, they were then placed in a fluctuating environment, and lastly, they were placed back to the static environment. We observed that a bistable trait evolved in the fluctuating environment to exhibit stochastic switching had a high likelihood to be maintained in the population in the subsequent static environment. By analyzing the simulation results, we found that the bistable trait evolved in the fluctuating environment had opposing characteristics from the one maintained in the subsequent static environment; while the former had a low-threshold switch to ensure that a small fraction of the population can be adaptive to the adversarial environment (i.e., to be used for stochastic switching), the latter had a higher-threshold switch to reduce gene expression variability (i.e., to be used to increase gene expression stability). Our results give evidence that such hidden switching capabilities can be established and maintained in a static environment. Interestingly, this is not because bistable switching has selective advantages under stabilizing selection, but rather it is because heightening the barrier between the stable states is an easier solution to increase the stability of gene expression with small mutational shifts than reverting back to a monostable gene expression system.

## Results

### Gene circuit model and evolutionary simulation

We simulated the evolution of a population of 1,000 asexual microorganisms in varying environmental conditions (See Methods for detailed description of the simulation procedure). We represented each individual organism in the population by a stochastic gene circuit model regulating the expression of master regulatory protein X. The gene circuit model for the expression of protein X is composed of four reaction processes: transcription; mRNA degradation; translation; and protein degradation. This model has four evolvable parameters: *a*; *b*; *K*_*d*_; and *n*, where *a* represents transcriptional efficiency, *b* represents translational efficiency (or protein burst size), *K*_*d*_ represents binding affinity (or the activation threshold), and *n* represents binding cooperativity. In this simulation, the individuals in the founding population were isogenic and had the identical, monostable gene circuit, which was based on a constitutive promoter without feedback loop (i.e., *n* = 0).

The selection of individuals in the population was based on the expression of X, and the circuitry was allowed to form a positive feedback loop, which could give rise to a bistable switch with right combinations of parameters. We had two different environments *E*_0_ and *E*_1_ with different selective pressures. While environment *E*_0_ favors lower expression levels of protein X, environment *E*_1_ favors higher expression levels considering metabolic costs for the protein production (i.e., stabilizing selection towards a given optimal expression level).

In the simulation, the population was evolved for 30,000 generations under different conditions. During the first 10,000 generations of evolution, the population was placed in *E*_1_. We refer the evolved population in this first environmental condition to as the *ancient population*. For the next 10,000 generations of evolution, the population was then placed in a fluctuating environment that switches environments between *E*_0_ and *E*_1_ every 20 generations. We call the evolved population in this second environmental condition the *intermediate population*. Finally, during the last 10,000 generations of evolution, the population was placed back in *E*_1_. We call the evolved population in this final environmental condition the *derived population*. We generated 50 sample evolutionary trajectories from this simulation procedure.

### Phenotypic characteristics determined by environmental conditions

From the simulation results, we first analyzed the fitness and the phenotypic characteristics of the three populations after they were evolved in their respective environments. For the evolved populations of each sample evolutionary trajectory, we measured the mean and the standard deviation of the fitness and the protein abundance level. Since the ancient and the derived populations (i.e., the evolved population from the first 10,000 generations and the evolved one from the last 10,000 generations) were evolved in the same static environment for a long period of generations, we expected many of the individuals in these populations to be adapted to exhibit highly optimized phenotypes for this environmental setting. Indeed, we observed that these two populations consistently evolved to have very high average fitness with low variances among individuals in each sample trajectory (Fig. 1a). By contrast, the intermediate population (i.e., the evolved population from the second 10,000 generations under the fluctuating environment) exhibited a widely different pattern with higher variability among the 50 trajectories, and in each of these trajectories, the population was often found to have higher fitness variances (Fig. 1a). Despite the higher variability, however, its average fitness was mostly on a par with or higher than the counterparts evolved in the static environment, suggesting that the intermediate population was able to adapt to *E*_1_ under our fluctuating selection.

**Figure 1.**
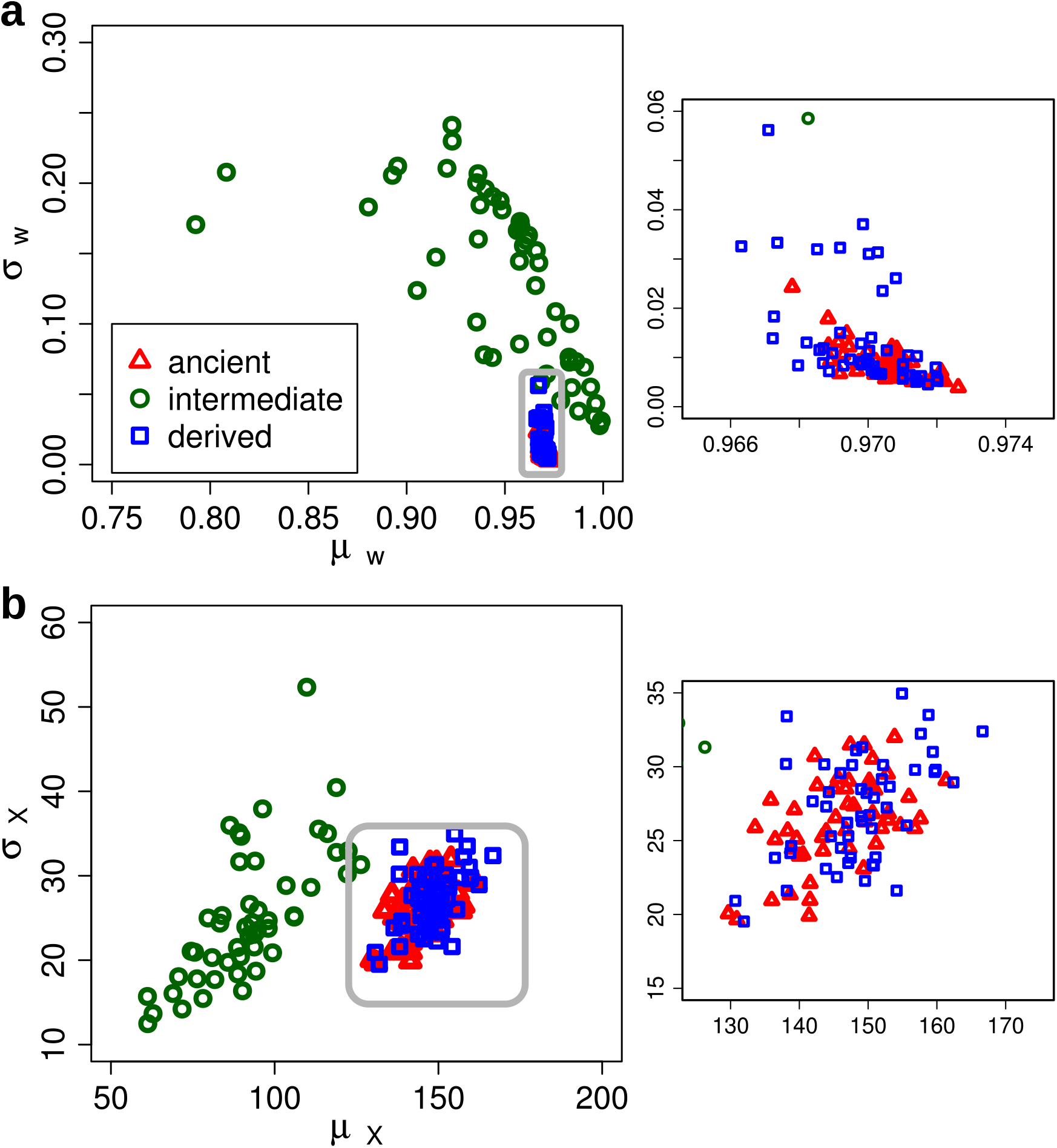
A comparison of phenotypic characteristics among the evolved ancient, intermediate, and derived populations for the 50 simulation runs. (a) A scatter plot showing the average fitness (*µ*_*W*_) and the standard deviation of the fitness (*σ*_*W*_) for each run. (b) A scatter plot showing the average protein level (*µ*_*X*_) and the standard deviation of the protein levels (*σ*_*X*_) for each run. In each plot, the right pane shows a close-up plot of the grey frame. The data were from evolved populations in the *E*_1_ environment: the 10,000th generation for the ancient and derived populations and the 9,900th generation for the intermediate population.

From the protein abundance distribution data, we also observed clear differences between the populations evolved in the static environment and the population evolved in the fluctuating environment (Fig. 1b). By comparing the protein abundance distributions, we found that the intermediate population frequently had relatively higher phenotypic variances given its lower average protein abundance levels than the derived population (Fig. 1b). This explains the high fitness variance, along with the lower average fitness, observed in the intermediate population. Such a high level of phenotypic variability among the individuals is advantageous in fluctuating environments, and it could arise from a heterogeneous population or a more homogeneous population with stochastic switching capability [14–21].

### Effects of the intermediate population on the evolution of genotypes in the derived population

The evolved individuals in the ancient and derived populations had similar phenotypes that were optimized for the static environment. There are two possibilities for the derived population to attain such phenotypic characteristics; it could either take the reversion course to have similar genotypes as the ancient population or evolve in a new direction to have novel genotypes optimized for *E*_1_. To discriminate these two possibilities, we compared the evolved parameters representing the most common genotype in the derived population against those in the ancient population and the intermediate population (Fig. 2). The comparison of the evolved parameters between the ancient and the derived populations revealed that they have weak correlations, indicating that the evolved genotypes in the derived population are widely different from those from the ancient population (Fig. 2a).

**Figure 2.**
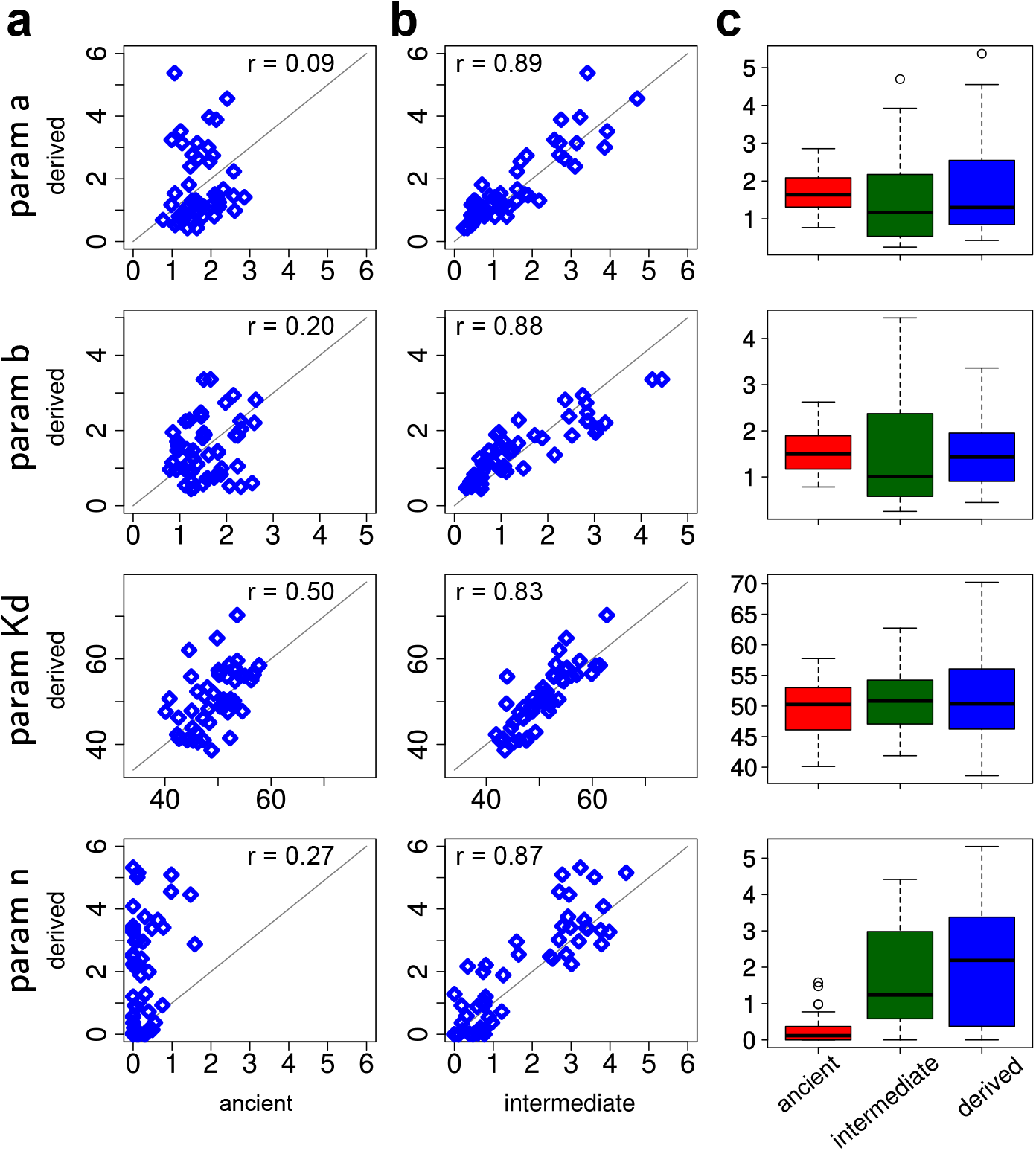
A comparison of genotypic characteristics among the ancient, intermediate, and derived populations for the 50 simulation runs after 10,000 generations of evolution. (a) Scatter plots showing the four evolved parameters for the most common genotype in the ancient and derived populations for each run. (b) Scatter plots showing the four evolved parameters for the most common genotype in the intermediate and derived populations for each run. (c) Box plots showing statistics of the most common genotype in the three evolved populations over the 50 runs.

What caused the genotypic differences between the ancient population and the derived population? The only difference between the two in our simulation setting is the initial genotypes in their populations; the derived population inherited genotypes from the intermediate population, whereas the ancient population began with a homogeneous genotype that exhibits a monostable gene expression. Thus, the genotypes evolved in the fluctuating environment must have played a crucial role in directing the derived population to take a different strategy and to form novel genotypes optimized for environment *E*_1_. Indeed, the evolved parameters in the derived population showed much stronger positive correlations with those in the intermediate population (Fig. 2b). The comparison of the evolved parameters using their average values also indicated similar relation patterns among the three populations (Supplementary figure S1).

To analyze how the genotypes of the intermediate population contributed to the evolution of novel genotypes in the derived population, we examined the distribution of each evolvable parameter in the three populations (Fig. 2c and Supplementary figure S1c). The clearest pattern that emerged from this was that the intermediate and derived populations tended to have high levels of binding cooperativity (i.e., high values of *n*) which indicate highly nonlinear transcription processes with strong positive feedback, whereas the ancient population had very low levels of binding cooperativity (i.e., low values of *n*).

### Maintenance of bistability under the stabilizing selection

Since nonlinear transcription with strong positive feedback can give rise to bistability and stochastic phenotypic switching [22], we set out to examine the shape of protein abundance level distributions for the evolved individuals and to classify whether their gene regulatory processes were monostable or bistable (see Methods). To this end, we analyzed the evolved parameter sets and counted the number of monostable and bistable individuals in the ancient, intermediate, and derived populations (Fig. 3). We found statistically significant differences in the characteristics of gene expression control between the ancient and the derived populations—the two populations that were evolved under the identical, static selection scheme (*p* < 10^*−*8^ with Fisher’s exact test). As expected, the bistable gene expression trait were not observed in the ancient population in all of the sample evolutionary trajectories. By contrast, the derived population repeatedly expressed the bistable trait (48%: 24 out of 50 sample evolutionary trajectories), and evolved individuals with the bistable trait often accounted for a large portion of the population. Indeed, we observed that more than 60% of sample evolutionary trajectories have the derived population with the bistable trait as the majority (15 out of 24).

**Figure 3.**
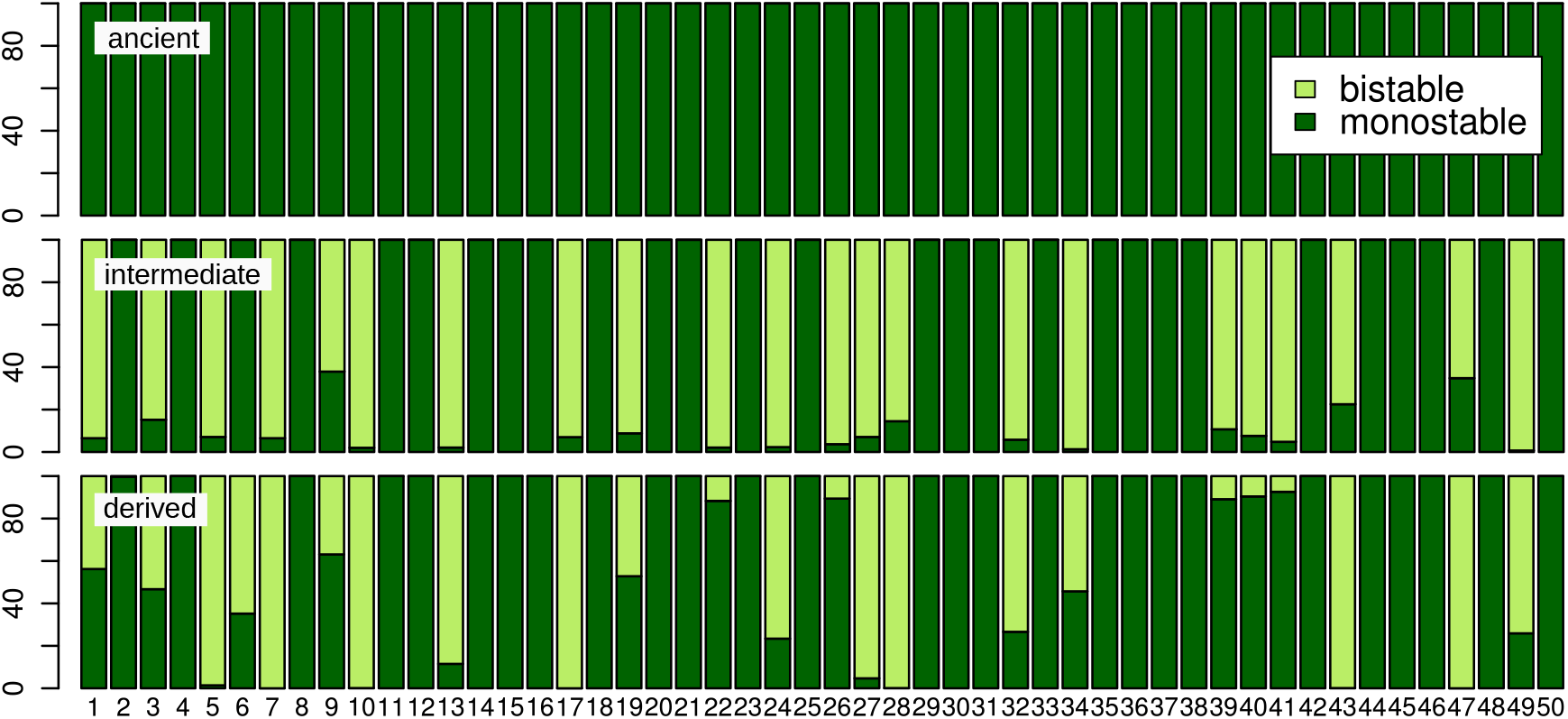
The percentage of the monostable and bistable individuals in the evolved ancient, intermediate, and derived populations for the 50 evolutionary simulation runs. The x-axis shows the run number of each simulation. A dark green region indicates the percentage of monostable individuals whereas a light green region indicates the percentage of bistable individuals in a given evolved population.

To understand the extent to which the intermediate population plays a role in the expression of the bistable trait in the derived population, we analyzed the relation between the gene expression traits of the intermediate population and the derived population. By computing the conditional probability, we found that when the bistable trait was evolved in the intermediate population, this trait was likely to be maintained in the derived population (100%: 22 out of 22). On the other hand, when the intermediate population only had the monostable trait, the derived population had a much lower chance to evolve the bistable trait (~ 7%: 2 out of 28). Our results indicate that the expression of the bistable trait in the derived population depends strongly on the evolution of the trait in the intermediate population.

To test if these results depend strongly on our specific choice of fitness function, we performed evolutionary simulation with a different type of fitness function that modeled stabilizing selection in the static environment (see Methods). We similarly observed that the derived population had a high propensity to express bistable individuals when the intermediate population expressed bistable individuals (Supplementary Figure S2), suggesting that these phenomena are independent of specific choices of fitness function. We further examined if our results depend on our specific choice of the size of mutational shifts. Because larger changes in the binding cooperativity can spontaneously turn monostable individuals into bistable ones and vice versa, we performed additional evolutionary simulations with different sizes of mutational shift for this parameter (see Methods). We found that, although larger mutational shifts in binding cooperativity resulted in the expression of the bistable trait in the ancient population, the fraction of the bistable trait observed in the derived population was much higher and that the the dependency of the bistable trait expression in the derived population on the intermediate population was strong (Supplementary Figures S3 and S4). From these additional experiments, We confirmed the consistency of our qualitative results under various evolutionary simulation settings.

### An increase in the stability of gene expression in static environment

But, how could the bistable trait evolved in the intermediate population be stably maintained in the derived population? Under fluctuating selection, bistability and stochastic phenotypic switching could evolve as a byproduct to increase evolvability [21]. Since stochastic switching allows a fraction of individuals to adapt to adversarial environments without genetic mutations, the population can increase the overall fitness rapidly [23–25]. Thus, we expected such a strategy be fixated in the intermediate population once evolved. In the static environment, however, random switching between the alternative phenotypes would result in unfit individuals without any obvious advantages in the population. That is, under stabilizing selection, more advantageous characters are those with an increase in the stability of gene expression levels to constantly express optimized phenotypes [26].

Thus, we suspected that the direction of evolution in the static environment was associated with an increase in the stability of gene expression process near the optimal gene expression level. One mechanism to lower gene expression noise (i.e., increase the gene expression stability) is to increase transcriptional efficiency and to decrease translational efficiency [14,27], and interestingly, we observed that the ancient population evolved higher transcriptional efficiency (i.e., higher levels of *a*) and lower translational efficiency (i.e., lower levels of *b*) once the gene expression rate reaches near the optimal level (Supplementary figure S6). Thus, this gives evidence to support the view that individuals with higher gene expression stability were selected for in the ancient population. We were not, however, able to detect similarly clear features for higher gene exrepssion stability in the derived population (Supplementary figure S6). This may be due to a complex relation between the gene regulation parameters and the dynamical behavior given higher levels of nonlinearity.

To better understand the relation between the gene expression stability and the evolution under stabilizing selection, we quantified the stability level of the gene expression process in the individuals in the ancient and the derived populations. To this end, we defined a stability measure around the higher expression stable state by assuming that the distribution of the protein abundance around the stable state be well approximated by a Gaussian function (see Methods for details). Using this stability measure, we computed the average evolution of the average gene expression stability of each population over the 50 sample trajectories (Fig. 4a). From this, we found that the ancient and the derived populations both similarly increased their gene expression stability levels as they evolved before they settled down at similar levels of gene expression stability. Since the initial stability level of the derived population was much lower than the final stability level of the ancient population, we can also deduce that the stability level decreased as the intermediate population evolved under fluctuating selection (Fig. 4a). Indeed, by comparing the average stability levels of the evolved individuals in the intermediate population and the derived population, we found that, in all of the 50 sample trajectories, the average stability level in the derived population was higher than the corresponding intermediate population (Fig. 4b). Thus, an increase in the stability level was a discriminative feature which was only observed in the evolution under stabilizing selection.

**Figure 4.**
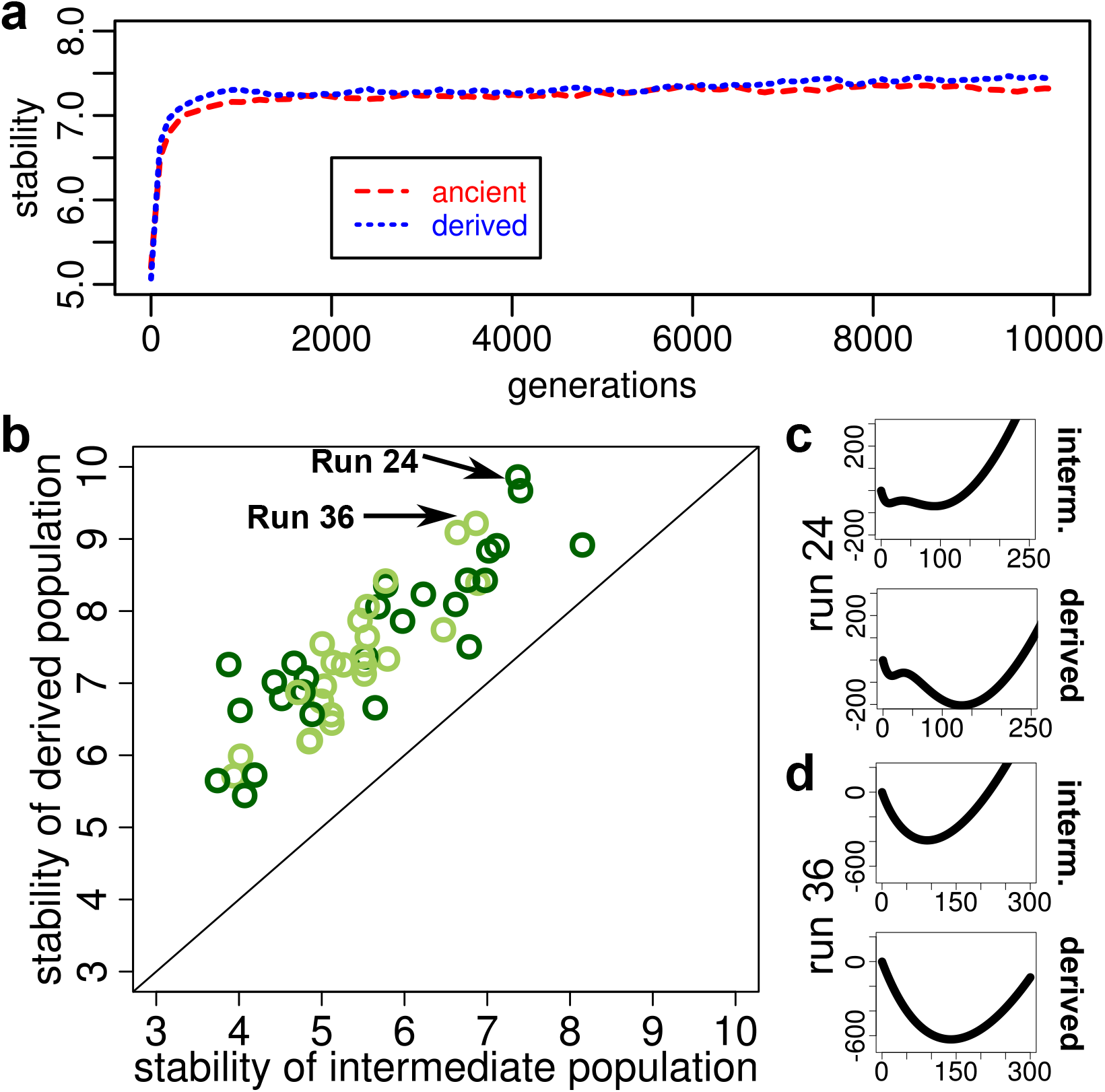
Evolution of gene expression stability. (a) The mean evolution of the average stability around the higher expression stable state between the ancient and derived populations over the 50 evolutionary simulation runs. (b) The average stability around the higher expression stable state between the evolved individuals in the intermediate and derived populations. Each dark green point represents a run with the derived population with at least one bistable individual, while each light green point represents a run with the derived population only consisting of monostable individuals. (c) A change in the potential function of the fittest individual in the intermediate and derived populations for a sample run producing a bistable derived population. (d) A change in the potential function of the fittest individual in the intermediate and derived populations for a sample run producing a monostable derived population.

### Maintenance of bistability as a general approach to increase the stability of gene expression

We examined the mechanism in which the gene expression stability was evolved in the static environment. To this end, we measured the potential landscape of the gene expression process of the fittest individuals from samples in which the derived population had high levels of average gene expression stability (see Methods). From this, we found that, although diverse monostable and bistable individuals evolved in the derived population, they shared a common phenotypic character that deepens the potential well around the stable state to increase the gene expression stability near the optimal gene expression level (Fig. 4c and d).

To test if the maintenance of the bistable trait to increase the gene expression stability is a general strategy that is independent of the specific structure of underlying gene circuit models, we used a more general model based on Gaussian distributions (see Supplementary Section S1 for detailed descriptions). This abstract model represents a monostable gene expression process using a Gaussian distribution and a bistable one using a mixture of two Gaussian distributions. Transitions between the monostable and the bistable processes are modeled using the Gaussian width. In evolutionary simulation, we fixed the stable state of each gene expression model to a level relatively close to the optimal one under stabilizing selection, and changes in the gene expression stability were indicated by changes in the value of the Gaussian width. We evolved clonal populations of monostable and bistable founders under various settings. In all simulation settings, we consistently observed that the Gaussian width gradually decreased (i.e., the gene expression stability gradually increased) as the population evolved (Fig. 5 and Supplementary Figures S5, S6, and S7). When the founding population consisted of monostable individuals, the monostable trait was maintained and lower Gaussian widths (i.e., those with higher gene expression stability) were selected for and maintained during the evolution. Similarly, when the founding population consisted of bistable individuals, the bistable trait was maintained and higher levels of gene expression stability were evolved during the evolution.

**Figure 5.**
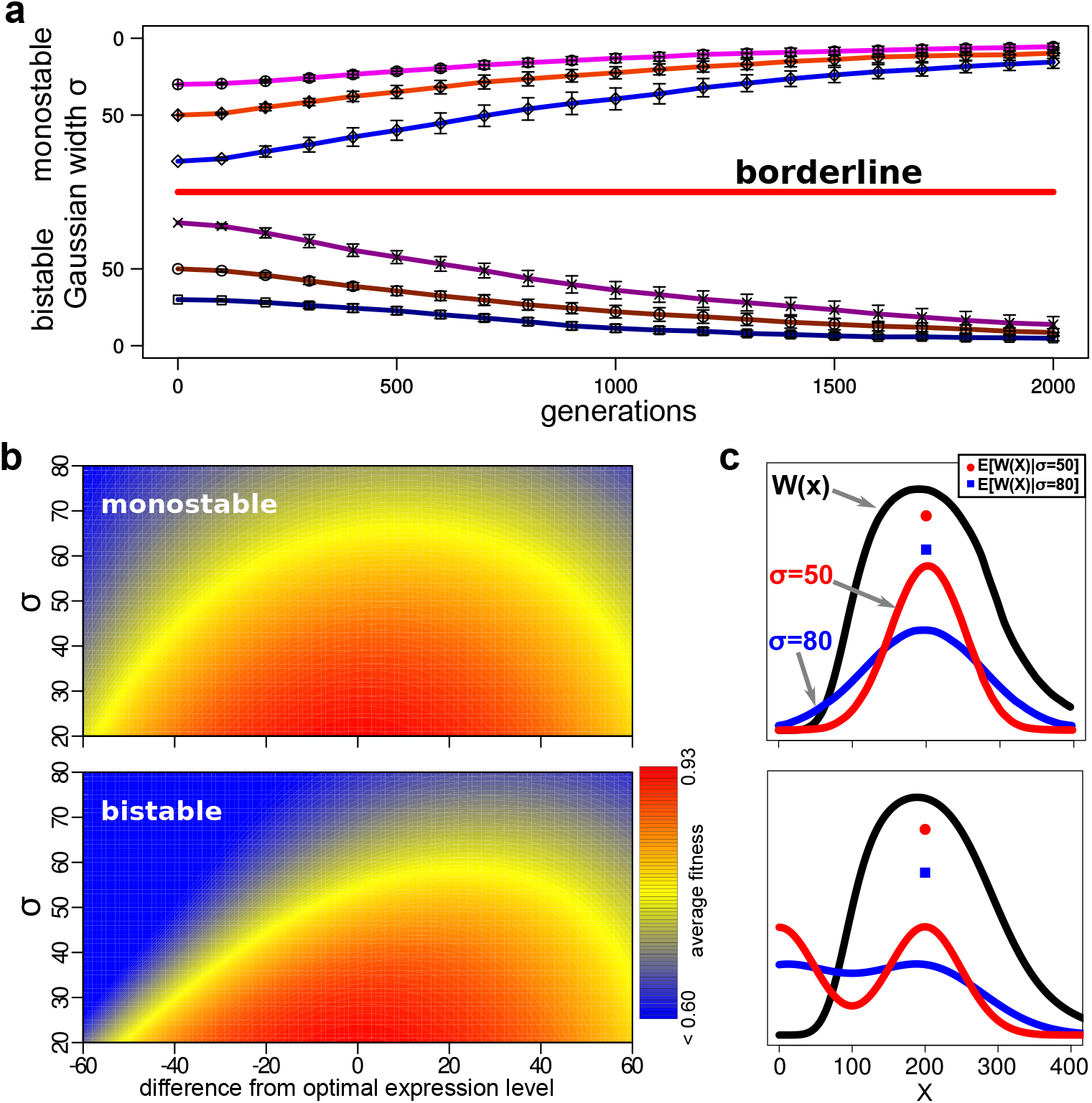
Results from evolutionary simulations of Gaussian-based gene expression models. (a) Average evolutionary trajectories of a Gaussian-based gene expression model. Evolution was simulated from a clonal population of either monostable individuals or bistable individuals. There were three different settings for the initial value of the Gaussian width (i.e., gene expression stability): 30, 50, and 80. 20 sample trajectories were generated from evolutionary simulations for each setting. Each point represents the average of the population average Gaussian width of the 20 runs, while each error bar represents the standard deviation of the population average. (b) Heat maps showing how the average fitness value changes based on the Gaussian width and the distance of the mean expression level from the optimal gene expression for the monostable model (top pane) and the bistable model (bottom pane). (c) Difference in the average fitness based on different value of the Gaussian width for the monostable model (top pane) and the bistable model (bottom pane).

## Discussions

Our results have shown that high levels of gene expression stability evolve under stabilizing selection and, to acquire such phenotypic characters, different evolutionary strategies are taken depending on the genotypic characteristics of the founding population. In a population consisting of monostable individuals, the evolution favored the monostable genotype that deepens the potential well around the stable state. By contrast, in a population with bistable individuals, the evolution was directed to raise the potential barrier that separates the bistable potential wells, increasing the stability around the high expression stable state. The reversion course from the bistable to the monostable was selected against under our evolutionary simulation with gradual mutational shifts because such a move would have inevitably decreased the gene expression stability temporarily in transition. As a way to acquire characters with high levels of gene expression stability, thus, keeping intricate molecular mechanisms to have bistability offered a better and simpler solution in such cases. These results demonstrate a plausible evolutionary scenario in which a wide-type population can stably maintain the switching machinery during a long period of environmental stasis. That is, low-threshold switches that are induced as a response to selective pressures in fluctuating selection can be gradually evolved to establish high-threshold switches that buffer gene expression variability to consistently produce expression levels adaptive to a given static environment.

There is evidence supporting that high levels of gene expression stability facilitated by hidden switching mechanisms play essential roles in development, which includes the normal workings of the yeast galactose-signalling network [28] and the *Xenopus* oocyte maturation decision network [29]. More recently, Raj, et al. [10] observed that some mutations in the wild-type *Caenorhabditis elegans* intestinal development circuit resulted in a highly stochastic cell-fate decision with some mutant embryos failing to develop intestinal cells. They found that the activation of *elt-2*, the master regulator of intestinal differentiation, via its feedback-based bistable regulation is crucial to the normal intestinal differentiation, and mutations in the upstream genes in this network can increase the expression variability in *elt-2*, leading to abnormal development. Our results suggest that some of such hidden switches might have been maintained to facilitate higher phenotypic stability under natural selection.

While our results indicated that high levels of gene expression stability is advantageous in static environments, the evolved stability levels clearly showed the presence of an upper bound. Because our gene circuit model captures the intrinsic fluctuations of gene expression processes along with other sources of the gene expression variability, the plateau of the gene expression stability that we observed in both the ancient and the derived populations may reflect on the fundamental limit to contain gene expression noise [30–32]. While the level of this limit depends on the specific structure of gene regulatory processes, an important constraint applicable to all biological systems is that some level of variability from the intrinsic fluctuations is always inevitable, and evolution would deal with such physical constraints. Our results suggest that an increase in the gene expression stability is advantageous when the mean expression level is close to the optimum and is positively selected for under natural selection. This does not contradict a previous study which considered the evolutionary scenario in which the average gene expression level is set to be fixed and far from the optimal level and concluded that higher levels of gene expression noise would be beneficial and selected for under stabilizing selection [33]. That is, our study was concerned with the evolution of gene expression processes that are allowed to adjust their mean expression level via advantageous mutational shifts in the model parameters. In our simulation, if the current gene expression level were to be far from the optimal one, the evolution could simply select mutants that move the gene expression level closer to the optimal level in the static environment. Thus, our focus was more on the evolution of gene expression stability when the mean expression level is close to the optimum. These differences can change the effects of gene expression stability substantially. Indeed, a recent experimental study in yeast confirmed that the effects of gene expression noise on fitness depends on the distance between the average protein abundance level and the optimal abundance level; higher noise levels are advantageous when the average protein level is far from the optimal one, whereas lower noise levels are advantageous when the average protein level is close to the optimum [34].

We found that genotypes to express the high-threshold bistable phenotype were very similar to those for the low-threshold one. Thus, “hidden” bistable switches stably maintained in a population to buffer gene expression variability in a static environment could be reactivated to drastically change the gene expression profile and release cryptic phenotypes via relatively small genetic drifts. Such phenotypic switching may play a crucial “capacitor” role in unveiling cryptic genetic variations and facilitating the evolution of adaptive novelties [7, 35]. In 1942, Waddington used the term canalization to describe general observations that naturally selected organisms produce one definite end-product regardless of minor variations in conditions during the development [1]. Although stochastic gene expression was not considered in his model of canalization [36], recent studies based on single-cell experimental methods revealed that the ability to control gene expression noise in wild type was essential to the constancy of developmental program and the complete penetrance of phenotypes [9, 10]. Thus, regulation of gene expression noise is crucial to the buffering of underlying variations and to the developmental canalization of discrete phenotypes. Our study shows that this extended view of canalization can explain the evolution of hidden threshold traits from stochastic switching traits and the stable maintenance of high-threshold bistable switches in naturally selected organisms in static environments.

## Methods

### Gene circuit model

Our gene circuit model for the expression of protein X is composed of two variables: *m* and *x* representing the molecule copy numbers of the mRNA and the protein forms, respectively, and four reaction processes: transcription; mRNA degradation; translation; and protein degradation. The transcription process is modeled using an thermodynamics-based approach and has the following kinetic law:

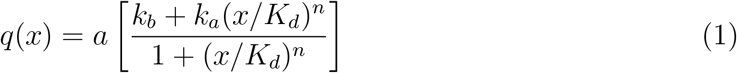

where parameters *k*_*b*_ and *k*_*a*_ are the basal and the activated transcription rates, respectively, with *a* being a scaling factor of these rates, while *K*_*d*_ and *n* represent the binding affinity and cooperativity, respectively.

The mRNA degradation process is modeled using a first-order reaction with kinetic law: *k*_*mdeg*_*m*, while the translation process is modeled using another first-order reaction with kinetic law: *k*_*trans*_*m*. The average number of the protein molecules produced from a single copy of mRNA transcript, then, is given by *b* = *k*_*trans*_/*k*_*mdeg*_. The protein degradation process is modeled using a first-order reaction with kinetic law: *k*_*deg*_*x*. To simulate this gene circuit model for one cell generation, we ran Gillespie’s stochastic simulation algorithm [37] for 2,000 seconds (around 30 minutes), which is within the range of typical bacterial generation time.

To map the genetic makeup of each individual with its phenotype, we needed an approach to obtain the dynamical property of the gene circuit model with a given parameter combination. With a continuum-state approximation, a stochastic process representing this gene circuit can be described by the following Fokker-Planck equation:

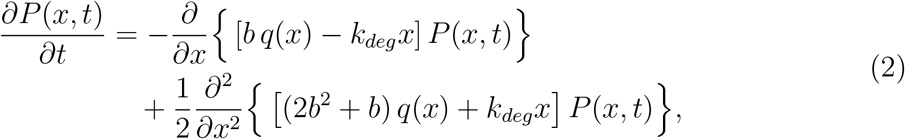

where *P* (*x, t*) is the probability that the protein level is *x* at time *t*. From this equation, the time-invariant probability distribution of the protein abundance level in the stationary regime, *P*_*s*_(*x*), is given by

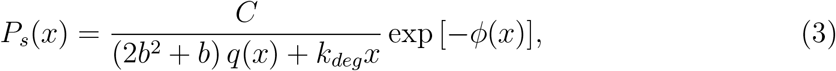

where

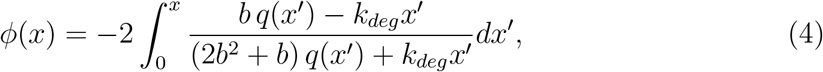

where *C* is a normalization constant and *φ*(*x*) is the potential function. By approximating *dx* by a small but finite ∆*x*, we can use this equation to map parameters to the protein abundance distribution without performing stochastic simulations.

### Evolutionary simulation

As described in Results, the evolution of the population of 1,000 individuals with this gene circuit was simulated for 10,000 generations under each of three different conditions consecutively. That is, during the first 10,000 generations, the population was evolved in environment *E*_1_, during the second 10,000 generations, it was evolved in a fluctuating condition under which the environment switches between *E*_0_ and *E*_1_ every 20 generations, and during the last 10,000 generations, the population was evolved in environment *E*_1_. In this simulation, we set *k*_*b*_ = 0.02, *k*_*a*_ = 0.2, *k*_*mdeg*_ = 0.1, *k*_*deg*_ = 0.002, while we set parameters, *a*, *b*, *K*_*d*_, and *n* as evolvable. The individuals in the founding population set these evolvable parameters as *a* = 1, *b* = 1 (i.e., *k*_*trans*_ = 0.1), *K*_*d*_ = 50, and *n* = 0, each having a gene expression process with a constitutive promoter without feedback loop regulation. Genetic mutations were captured by adding small perturbations to the evolvable parameters. Specifically, perturbations in *a*, *b*, *K*_*d*_, and *n* were first modeled using random variates from zero-mean Gaussian distributions with *σ* being 0.2, 0.2, 1, and 0.2, respectively. In our followup simulations, we have changed this perturbation setting to check the consistency of our results (see Supplementary figures S3 and S4). A mutation was introduced randomly to each individual with the rate of 0.01 per gene circuit per generation. By assuming that the gene circuit is of size around 1Kb, we can show that the population times the mutation rate per base pair per generation is 0.01, which is of the same order of magnitude and comparable with what has been reported [38] as well as those based on estimated effective population sizes (i.e., *N*_*e*_ ranging from 2.5 × 10^7^ to over 10^8^ [39, 40]) and estimated mutation rates per base pair per generation (*µ* ranging from 2.6 × 10^−10^ to 5.4 × 10^−10^ [40, 41]) of prokaryotes.

The fitness of individuals depends on their environment, and we have two fitness functions *W*_0_ and *W*_1_ for environments *E*_0_ and *E*_1_, respectively. These functions have the following forms:

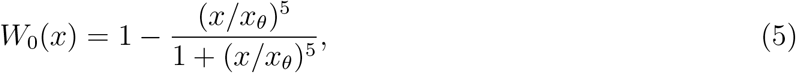

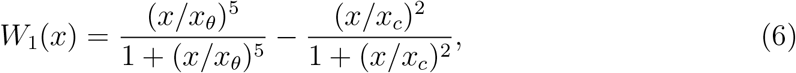

where *x*_*θ*_ is set to 50 and *x*_*c*_ is set to 1000. In environment *E*_0_, the optimal protein level, *µ*_*low*_, is 0, while in environment *E*_1_, the optimal level, *µ*_*high*_, is 135.

To test if our main results could be reproduced independent of specific fitness function we used, we performed evolutionary simulations with following fitness functions:

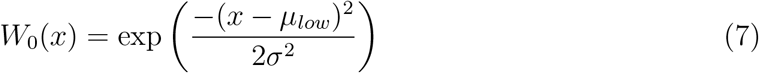

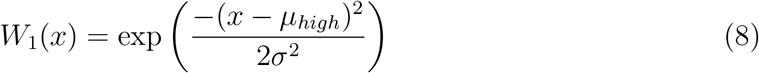

where *µ*_*low*_ (the optimal gene expression level for *E*_0_), *µ*_*high*_ (the optimal gene expression level for *E*_1_), and *σ* were set to 0, 150, and 85, respectively.

Selection of individuals for a new generation was modeled using roulette wheel selection. That is, to select each of 1,000 individuals for the new population, the normalized fitness over all individuals in the current population was used as the probability distribution for the selection, and each individual for the new generation was randomly picked from this distribution.

### The classification of bistability and monostability

To classify each individual in the population as a bistable or monostable gene expression phenotype, we noted that the sufficient condition to have either a stable state (i.e., protein level at which *P*_*s*_(*x*) has a peak) or an unstable state (i.e., protein level at which *P*_*s*_(*x*) has a bottom) is 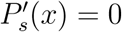. The first derivative of *P*_*s*_(*x*) can be expressed as

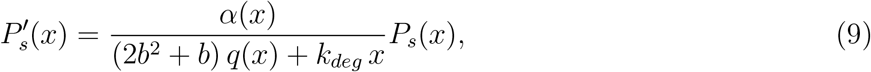

where

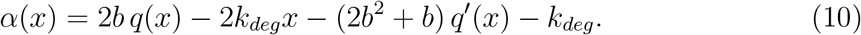

From this equation, we can see that stable and unstable steady states must be roots of *α*(*x*). We used a root-finding method to find the number of steady states and to classify whether a given genotype encodes a monostable gene expression process or a bistable gene expression process. With the boundary between 0 and 1000, we first count the number of roots of *α*(*x*) for each individual. Since we are interested in “functional” phenotypic switches, we considered only those bistable systems with a higher expression stable state being higher than threshold *x*_*θ*_ as bistable. Thus, if an individual has 2 roots of *α*(*x*) with the higher one being larger than *x*_*θ*_ (i.e., the higher expression stable state is higher than *x*_*θ*_ and the lower expression attractor state is 0), then we consider this as bistable. Also, if an individual has at least 3 roots with the 3rd smallest root being larger than *x*_*θ*_ (i.e., the higher expression stable state is higher than *x*_*θ*_), then we consider this as bistable. If a given individual does not satisfy either of these 2 bistable conditions, then we consider this individual as monostable.

### Stability measure

To quantify the stability of the individuals in the ancient and the derived populations, we approximated the shape of *P*_*s*_(*x*) around the higher expression stable state *x*_*hs*_ by a Gaussian function *g*(*x*) [32] where

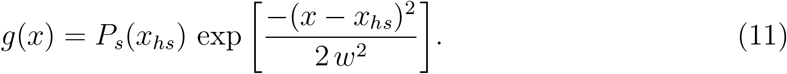

To obtain the value of width *w*, we set a constraint which requires 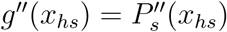, that is, the concavity around the higher expression stable state is set to be the same between the two functions. With this constraint, we can express *w* as

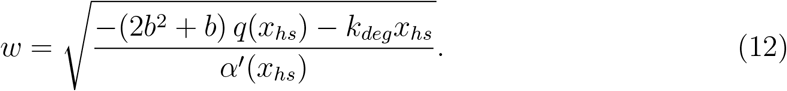

For each individual with *x*_*hs*_ > *x*_*θ*_, then, the stability of its gene expression process for the environment under which higher expression of protein X is favored is given by *x*_*hs*_/*w*. Thus, the higher this value is, the higher the gene expression stability around the higher expression stable state.

## Acknowledgment

We thank Orkun Soyer and Takashi Gojobori for their comments on an earlier version of the manuscript.

