## supplemental materials for "Stable maintenance of a hidden switch as a way to increase the gene expression stability"

### Supplementary Material for Stable maintenance of a hidden switch as a way to increase the gene expression stability

### Contents

**S1 Evolution with a Gaussian-based gene expression model** **3**

### S1 Evolution with a Gaussian-based gene expression model

In this follow-up evolutionary study, we were interested in analyzing if the maintenance of bistable traits was specific to the autoregulation gene expression model that we used or if the phenomenon was more general. To this end, we set out to use a gene expression model with a higher level of abstraction. In this abstracted model, we had two discrete-time stationary processes, $X_M(g)$ and $X_B(g)$, to represent the protein abundance levels of an monostable individual and a bistable individual at generation $g$, respectively. The probability density function of $X_M$ is denoted by $f_M(x)$, which is defined as follows:

$$f_M(x) = \begin{cases} \frac{C}{\sigma\sqrt{2\pi}} e^{-\frac{(x-\mu_H)^2}{2\sigma^2}}, & \text{if } x \geq 0, \\ 0, & \text{otherwise,} \end{cases} \tag{S1}$$

where $C = 1/\int_0^\infty (1/\sqrt{2\pi})e^{-(x-\mu_H)^2/2\sigma^2}dx$ is the normalization constant, $\sigma$ is the Gaussian width controlling the gene expression noise, and $\mu_H$ is the higher-expression stable state.

On the other hand, the probability density function of $X_B$ is bimodal which is based on two Gaussian distributions $f_L(x)$ and $f_H(x)$. If the protein abundance level at the previous generation, $x'$, is less than $x_\theta$, then the distribution $f_L(x)$ is used to sample $X$. Otherwise (i.e., if $x' \geq x_\theta$), then, the distribution $f_H(x)$ is used to sample $X$. That is, $f_L$ is the distribution for the lower expression valley and $f_H$ is the distribution for the higher expression one. Since the optimal expression level in the lower expression valley is 0, we set $f_L(x)$ to be the following half-normal density:

$$f_L(x) = \begin{cases} \frac{\sqrt{2}}{\sigma\sqrt{\pi}} e^{-\frac{x^2}{2\sigma^2}}, & \text{if } x \geq 0, \\ 0, & \text{otherwise.} \end{cases} \tag{S2}$$

In the higher expression region, the optimal expression level is the same as the monostable case, so we set $f_H(x)$ to be the following Gaussian density function:

$$f_H(x) = \begin{cases} \frac{C}{\sigma\sqrt{2\pi}} e^{-\frac{(x-\mu_H)^2}{2\sigma^2}}, & \text{if } x \geq 0, \\ 0, & \text{otherwise.} \end{cases} \tag{S3}$$

Thus, by using these two density functions, we defined $f_B(x \mid x')$, the density function for $X_B(g+1) = x$ given that $X_B(g) = x'$ as follows:

$$f_B(x \mid x') = \begin{cases} f_L(x), & \text{if } x' < x_\theta, \\ f_H(x), & \text{otherwise.} \end{cases} \tag{S4}$$

With this definition of the bistable probability distribution, if the protein level at the previous generation is at least $x_\theta$, the two probability density functions, $f_M(x)$ and $f_B(x \mid$

$x'$), become identical for this generation. On the contrary, if the protein level at the previous generation is lower than the threshold, the half Gaussian distribution with the lower-expression stable state (i.e., $f_L(x)$) is used. Thus, $f_B(x \mid x')$ has two Gaussian distributions separated by a barrier at $x = x_\theta$. Let $L$ and $H$ be the lower and higher expression regions, respectively, that is, $L = [0, x_\theta)$ and $H = [x_\theta, \infty)$. Based on this, the probability to spontaneously transition from $H$ to $L$ is given by

$$p_{H \to L} = \int_0^{x_\theta} f_H(x)dx, \tag{S5}$$

while the probability to spontaneously transition from $L$ to $H$ is given by

$$p_{L \to H} = \int_{x_\theta}^{\infty} f_L(x)dx. \tag{S6}$$

In the simulation reported in the main text (Fig. 5a), we set the value of $\mu_H$ and $x_\theta$ to 200 and 100, respectively. We also set different values for $\mu_H$ to see our conclusion still holds (see Supplementary Figure S6).

In this gene expression model, the only evolvable parameter was the Gaussian width $\sigma$. We captured transitions between the monostable trait and the bistable trait using the evolvable parameter $\sigma$. Specifically, we set the threshold for the Gaussian width $\sigma_\theta$ so that whenever $\sigma$ became higher than this threshold, we switched the trait to the other one. We set the value of $\sigma_\theta$ to 100 so that transitions occur when the potential function becomes flat—that is, when the gene expression stability becomes very low. A mutation in $\sigma$ was modeled by adding $\Xi_\sigma$, a zero-mean Gaussian random variable, to $\sigma$. To have gradual changes in the level of gene expression stability, the standard deviation of $\Xi_\sigma$ was set to 2. By denoting the Gaussian width of the $i$-th individual at generation $g$ by $\sigma_i(g)$, we modeled the change via mutation on $\sigma_i(g+1)$ as follows:

$$\sigma_i(g+1) = \begin{cases} \sigma_i(g) + \Xi_\sigma, & \text{if } \Xi_\sigma \leq \sigma_\theta - \sigma_i(g), \\ 0, & \text{if } \Xi_\sigma > -\sigma_i(g), \\ 2\sigma_\theta - \sigma_i(g) - \Xi_\sigma, & \text{otherwise.} \end{cases} \tag{S7}$$

In this simulation, the environment was assumed to be static and to always favor individuals with higher protein levels. To compute the fitness of each individual, we used the following function

$$W(x) = \frac{(x/x_\theta)^h}{1 + (x/x_\theta)^h} - \frac{(x/x_c)^8}{1 + (x/x_c)^8} \tag{S8}$$

where $h$ was set to 5 in the simulation reported in the main text, and in the follow-up simulation it was also set to 1 (see Supplementary Figure S4). With $h = 5$, the optimal gene expression level was around 189.2, which is close to $\mu_H$. We further performed simulations with a different form of this fitness function. Specifically, we used the following Gaussian-based fitness function

$$W(x) = e^{-\frac{(x-m)^2}{2s^2}}, \tag{S9}$$

where $m$ and $s$ were changed to take several values (see Supplementary Figures S5 and S6).

The number of individual in the population was fixed to be 1,000, and the mutation rate of the Gaussian width per individual per generation was set to be 0.01. In this evolutionary simulation, we used the same approach for the selection of individuals in the next generation based on the relative fitness.

### Figures

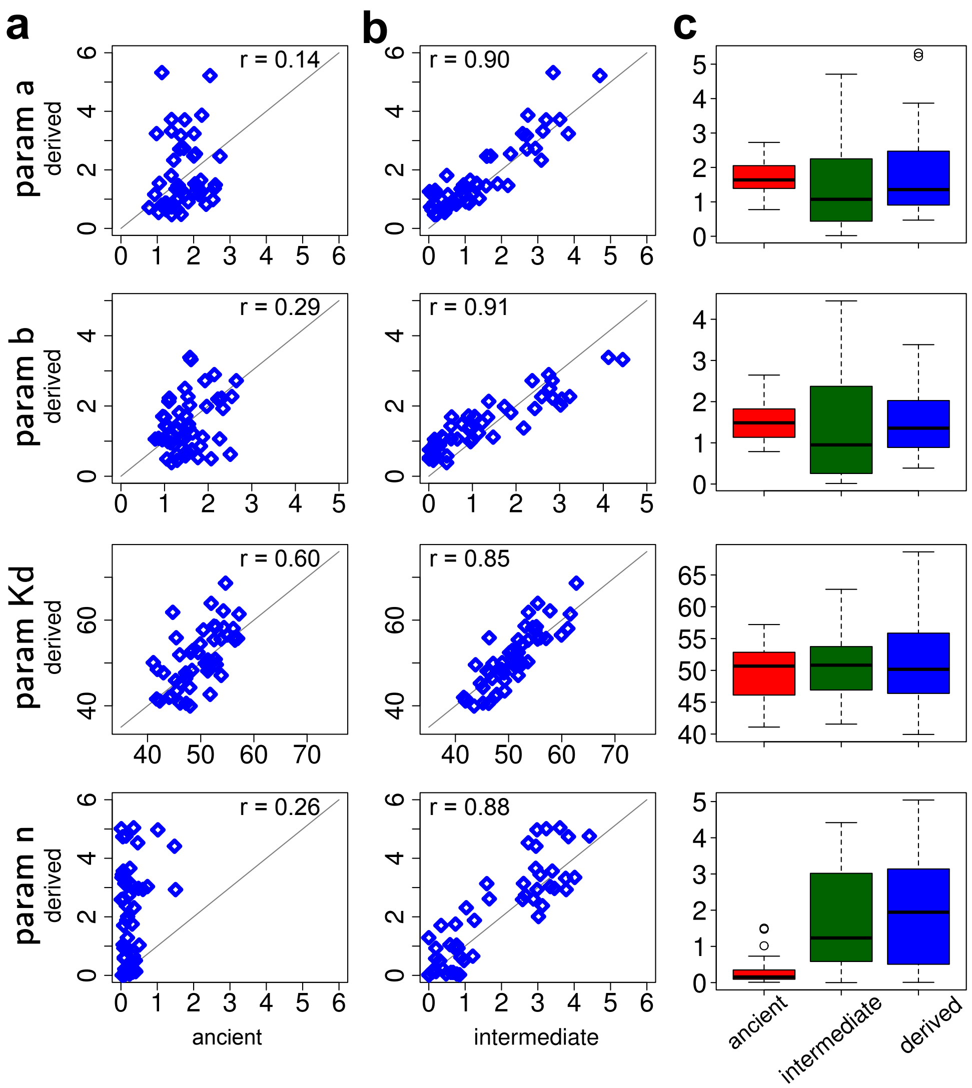


**Figure S1.** Evolution of the average mutable parameters in the derived population over the first 5,000 generations. Each point represents the average value of a parameter in the last generation of the intermediate population (x-axis) and that of the derived population at a specified generation (y-axis).

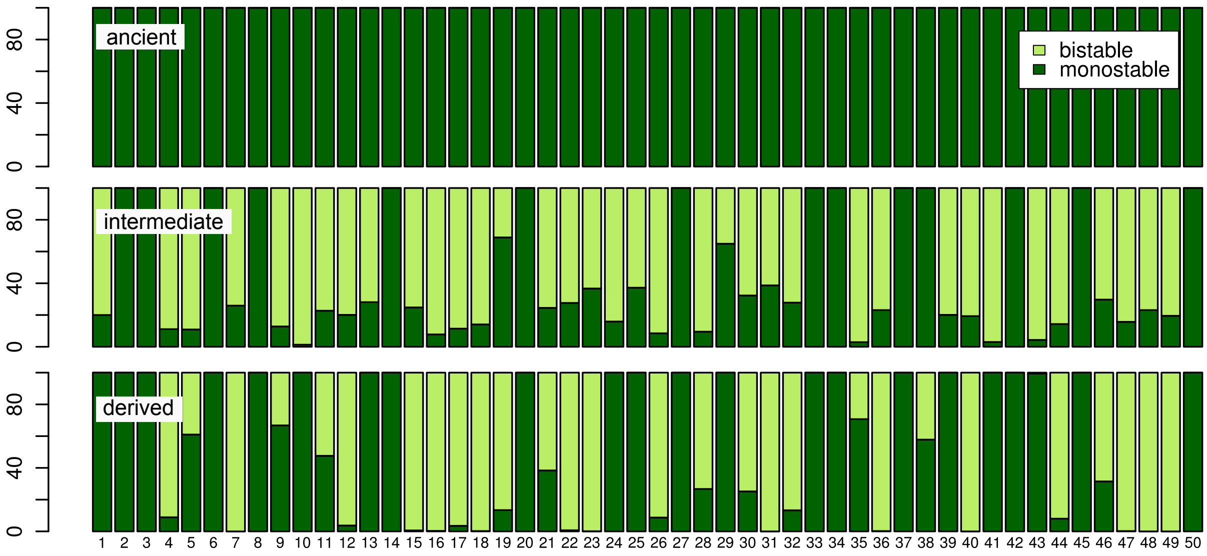


**Figure S2.** The proportion of monostable and bistable individuals in the 3 types of populations for the 50 evolutionary simulations. The fitness function used here is based on a Gaussian function, which is decribed in the Methods section in the main text. For each population, the data from the last generation (10,000 generation) were used.

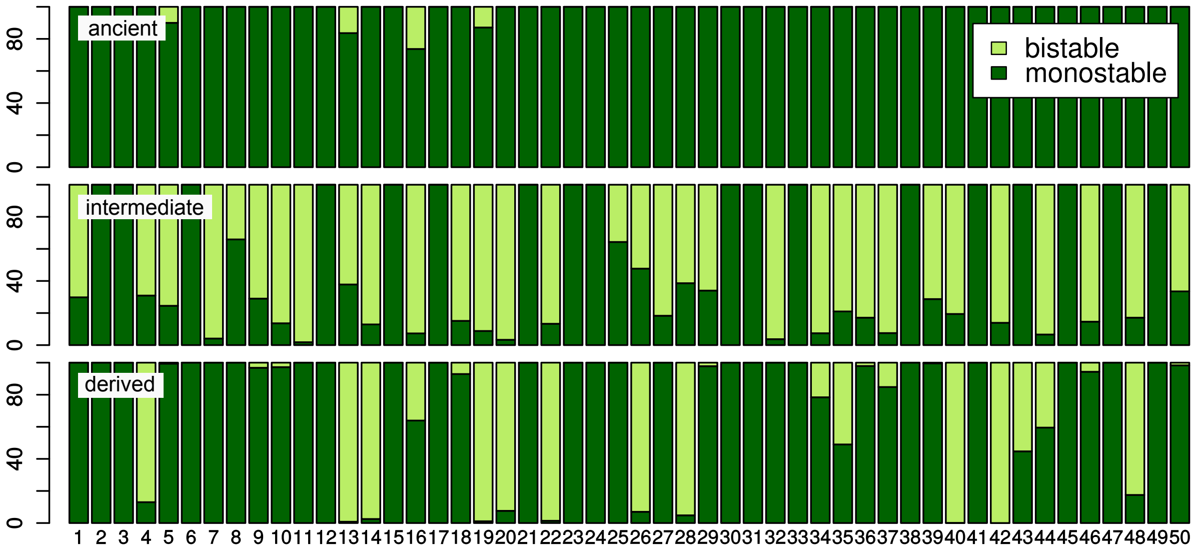


**Figure S3.** The proportion of monostable and bistable individuals in the 3 types of populations for the 50 evolutionary simulations. The mutational shift size of $n$ is changed to 0.5. For each population, the data from the last generation (10,000 generation) were used.

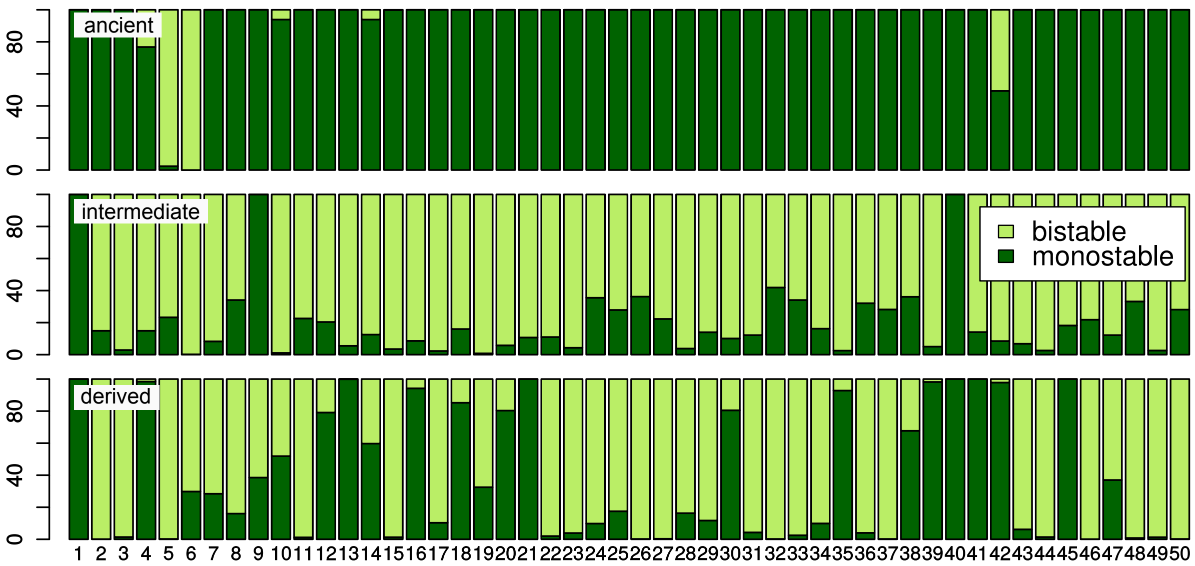


**Figure S4.** The proportion of monostable and bistable individuals in the 3 types of populations for the 50 evolutionary simulations. The mutational shift size of $n$ is changed to 0.6. For each population, the data from the last generation (10,000 generation) were used.

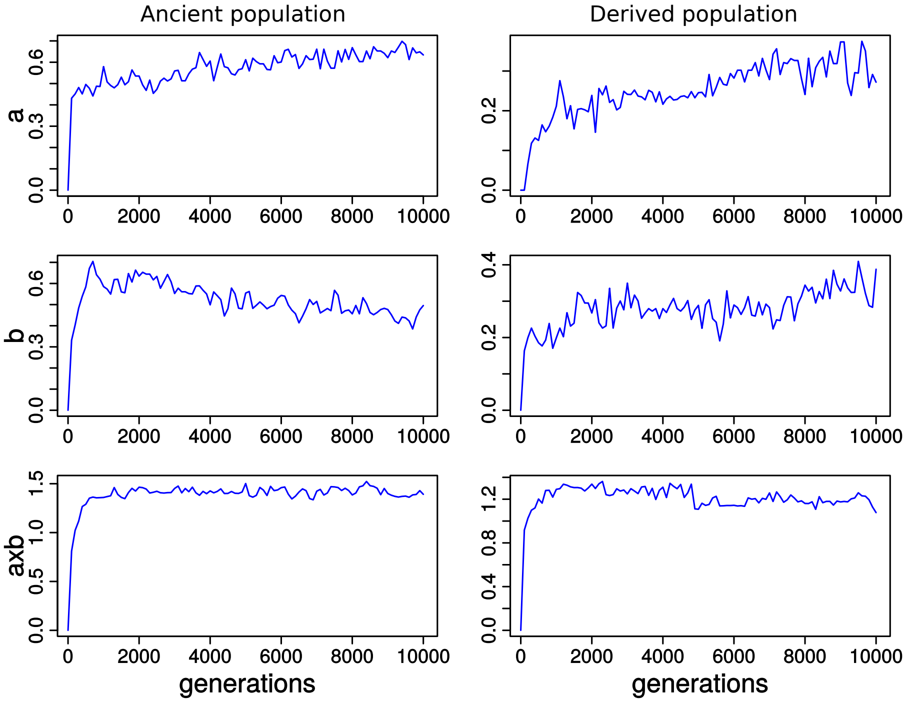


**Figure S5.** The relative change of each mutable parameter over generations in the derived population. The evolution of the median of the relative change of each mutable parameter in the derived population is shown with respect to the average value in the last generation of the intermediate population. The median value was taken from the population average of the 50 evolutionary simulation runs.

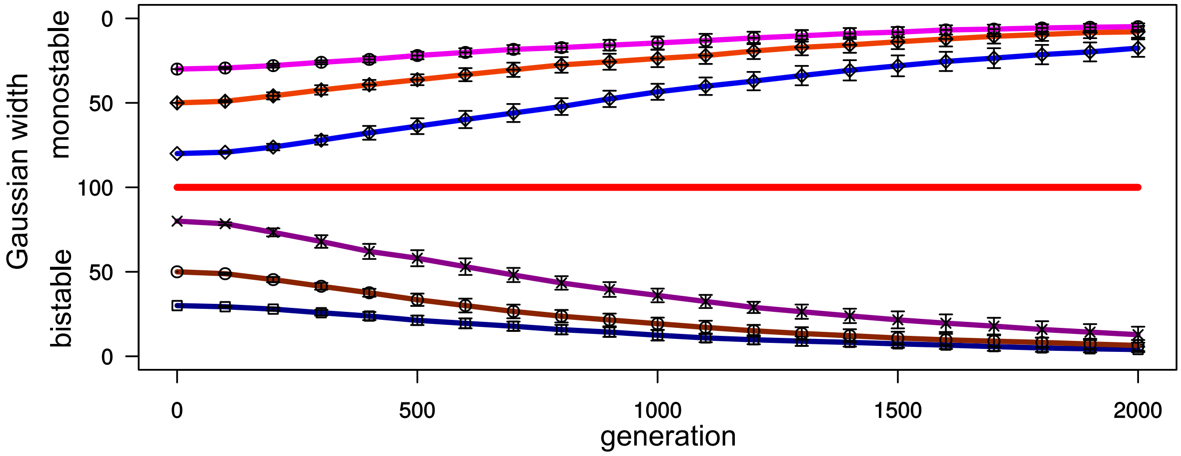


**Figure S6.** Average evolutionary trajectories of a Gaussian-based gene expression model based on a different type of fitness function. Here, $(x/100)/[1 + (x/100)] - (x/300)^8/[1 + (x/300)^8]$ was used as the fitness function. Evolution was simulated from a clonal population of either monostable individuals or bistable individuals. There were three different settings for the initial value of the Gaussian width (i.e., gene expression stability): 30, 50, and 80. 20 sample trajectories were generated from evolutionary simulations. Each point represents the average Gaussian width of the 20 runs, while each error bar represents the standard deviation.

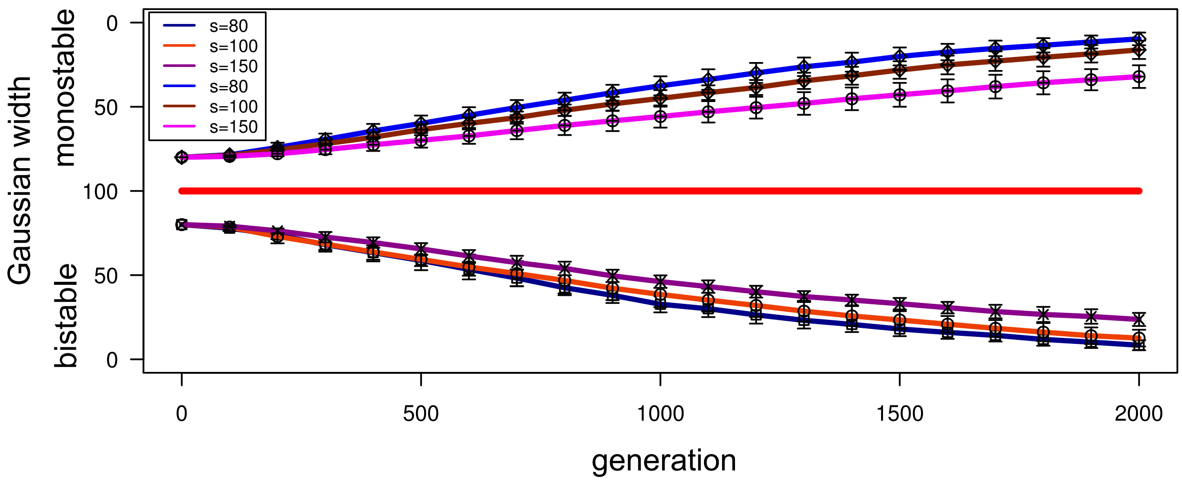


**Figure S7.** Average evolutionary trajectories of a Gaussian-based gene expression model with a Gaussian-based fitness function with various bell shapes. Here, the fitness function was changed to a Gaussian function $\exp\left[-(x-m)^2/2s^2\right]$ where $m$ was set to 200 and $s$ was decided to vary and take a value from 80, 100, and 150. The initial value of $\sigma$ for each setting was 80. Evolution was simulated from a clonal population of either monostable individuals or bistable individuals. There were three different settings for the initial value of the Gaussian width (i.e., gene expression stability): 30, 50, and 80. 20 sample trajectories were generated from evolutionary simulations. Each point represents the average Gaussian width of the 20 runs, while each error bar represents the standard deviation.

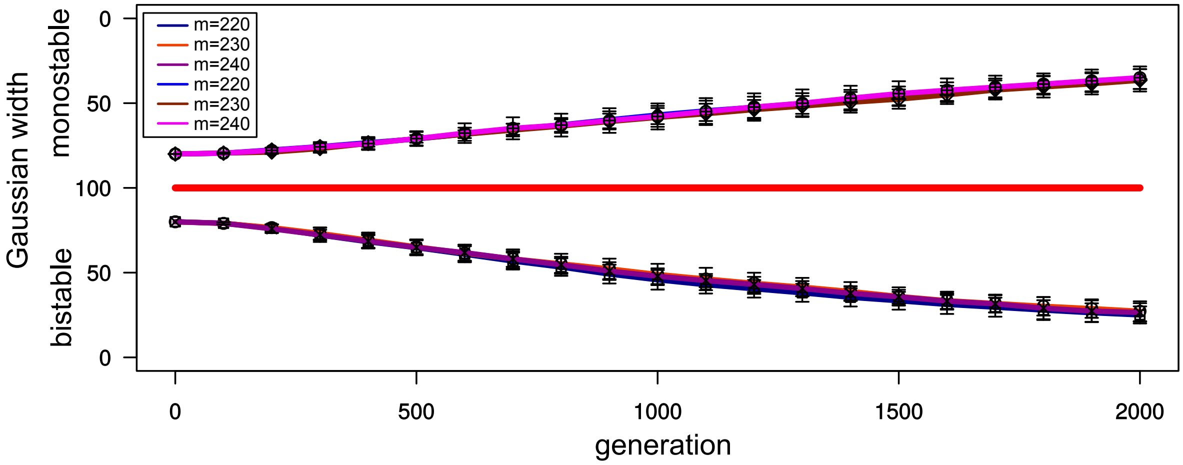


**Figure S8.** Average evolutionary trajectories of a Gaussian-based gene expression model based on various optimal high-expression level. A high-expression level is randomly chosen from a Gaussian function $\exp\left[-(x-m)^2/2s^2\right]$ where $s$ was set to 150 and $m$ was decided to vary and take a value from 220, 230, and 240. The initial value of $\sigma$ for each setting was 80. Evolution was simulated from a clonal population of either monostable individuals or bistable individuals. There were three different settings for the initial value of the Gaussian width (i.e., gene expression stability): 30, 50, and 80. 20 sample trajectories were generated from evolutionary simulations. Each point represents the average Gaussian width of the 20 runs, while each error bar represents the standard deviation.
